# Swine ANP32A supports avian influenza virus polymerase

**DOI:** 10.1101/2020.01.24.916916

**Authors:** Thomas P. Peacock, Olivia C. Swann, Ecco Staller, P. Brian Leung, Daniel H. Goldhill, Hongbo Zhou, Jason S. Long, Wendy S. Barclay

## Abstract

Avian influenza viruses occasionally infect and adapt to mammals, including humans. Swine are often described as ‘mixing vessels’, being susceptible to both avian and human origin viruses, which allows the emergence of novel reassortants, such as the precursor to the 2009 H1N1 pandemic. ANP32 proteins are host factors that act as influenza virus polymerase cofactors. In this study we describe how swine ANP32A, uniquely among the mammalian ANP32 proteins tested, supports some, albeit limited, activity of avian origin influenza virus polymerases. We further show that after the swine-origin influenza virus emerged in humans and caused the 2009 pandemic it evolved polymerase gene mutations that enabled it to more efficiently use human ANP32 proteins. We map the super pro-viral activity of swine ANP32A to a pair of amino acids, 106 and 156, in the LRR and central domains and show these mutations enhance binding to influenza virus trimeric polymerase. These findings help elucidate the molecular basis for the ‘mixing vessel’ trait of swine and further our understanding of the evolution and ecology of viruses in this host.

## Introduction

Influenza A viruses continuously circulate in their natural reservoir of wild aquatic and sea birds. Occasionally, avian influenza viruses infect mammalian hosts, but these zoonotic viruses have to adapt for efficient replication and further transmission. This limits the emergence of novel endemic strains. Avian-origin, mammalian-adapted influenza viruses have been isolated from a range of mammalian species including humans, swine, horses, dogs, seals, and bats (1–6).

One mammalian influenza host of significance are swine, which have been described as susceptible to viruses of both human- and avian-origin (6). It has been hypothesised that swine act as ‘mixing vessels’, allowing efficient gene transfer between avian- and mammalian-adapted viruses. This leads to reassortants, which are able to replicate in humans, but to which populations have no protective antibody responses, as best illustrated by the 2009 H1N1 pandemic (pH1N1) (7). The ability of pigs to act as ‘mixing vessels’ has generally been attributed to the diversity of sialic acids, the receptors for influenza, found in pigs that would enable co-infection of a single host by diverse influenza strains (8, 9). The husbandry of swine has also been hypothesised to play a role in this ‘mixing vessel’ trait; swine are often exposed to wild birds and it is likely their environments are often contaminated with wild bird droppings containing avian influenza viruses.

For an avian-origin influenza virus to efficiently infect and transmit between mammals several host barriers must be overcome. One major barrier is the weak activity of avian influenza virus polymerases in the mammalian cell. The acidic (leucine-rich) nuclear phosphoproteins of 32 kilodaltons (ANP32) proteins are key host factors responsible for the restricted polymerase activity of avian influenza viruses in mammalian cells (10). ANP32 proteins possess an N-terminal domain composed of five leucine rich repeats (LRRs) and a C-terminal low complexity acidic region (LCAR) separated by a short region termed the ‘central domain’. In birds and most mammals three ANP32 paralogues are found: ANP32A, ANP32B and ANP32E (11, 12). The roles of ANP32 proteins in cells are diverse and often redundant between the family members but include histone chaperoning, transcriptional regulation, regulation of nuclear export and apoptosis (12). In birds, such as chickens and ducks, an exon duplication allows for the expression of an alternatively spliced, longer isoform of ANP32A that effectively supports activity of polymerases of avian influenza viruses (10, 13). Mammals only express the shorter forms of ANP32 proteins which do not efficiently support avian polymerase unless the virus acquires adaptive mutations, particularly in the PB2 polymerase subunit, such as E627K (10). A further difference between the ANP32 proteins of different species is the level of redundancy in their ability to support influenza polymerase. In humans, two paralogues – ANP32A and ANP32B – are essential but redundant influenza polymerase cofactors (14, 15). In birds, only a single family member – ANP32A - supports influenza virus polymerase activity, as avian ANP32B proteins are not orthologous to mammalian ANP32B (11, 15, 16). In mice, only ANP32B can support influenza A polymerase activity (14, 15). Neither avian nor mammalian ANP32E proteins have been shown to support influenza polymerase activity (14–16).

In this study, we investigated the ability of a variety of mammalian ANP32 proteins to support influenza virus polymerases derived from viruses isolated from a range of hosts. We find differences in pro-viral efficiency that do not always coincide with the natural virus-host relationship: for example, human ANP32B is better able to support bat influenza polymerases than either bat ANP32 protein. Conversely, we describe evidence of human ANP32 adaptation early during the emergence of the pH1N1 virus from pigs, and find that swine ANP32A is the most potent pro-viral mammalian ANP32 protein tested, capable even of modestly supporting avian virus polymerases. This can be attributed to amino acid differences in the LRR4 and central domains that enhance the interaction between swine ANP32A and the influenza polymerase complex, suggesting a mechanism for this super pro-viral activity and giving support to the special status as potential ‘mixing vessels’ of swine in influenza evolution.

## Results

### Mammals naturally susceptible to influenza have two pro-viral ANP32 proteins

To investigate the ability of different mammalian ANP32A and ANP32B proteins to support influenza virus polymerase activity, several mammalian-origin influenza virus polymerase constellations were tested using an ANP32 reconstitution minigenome assay. A previously described human cell line with both ANP32A and ANP32B ablated (eHAP dKO) was transfected with ANP32A or ANP32B expressing plasmids from chicken, human, swine, horse, dog, seal or bat, as well as the minimal set of influenza polymerase expression plasmids for PB2, PB1, PA and nucleoprotein (NP), to drive amplification and expression of a firefly-luciferase viral-like reporter RNA and a *Renilla*-luciferase expression plasmid as a transfection control.

Initially, we tested a panel of polymerases derived from human, canine, equine and bat influenza viruses. In contrast to chicken ANP32B, which does not support influenza virus polymerase activity (11, 15, 16), chicken ANP32A and all mammalian ANP32A and ANP32B proteins supported activity of the mammalian-origin viral polymerases to varying degrees (Fig. 1a). Among the mammalian ANP32 proteins tested, for most polymerases, swine ANP32A provided the strongest support of polymerase activity, whereas the ANP32B proteins from dog, seal and bat displayed the least efficient pro-viral activity. These trends could not be explained by differences in expression levels or nuclear localisation (Fig. 1b, c). The bat influenza polymerases, along with (human) influenza B polymerase showed a different pattern of ANP32 usage, being able to strongly utilise ANP32Bs from all mammalian species, particularly human ANP32B (Fig. 1a). There was no evidence that influenza viruses adapted to particular mammals had evolved to specifically use the corresponding ANP32 proteins. For example, dog ANP32A or ANP32B were not the most efficient cofactors for canine influenza virus polymerase and human ANP32B was better able to support the bat influenza polymerase than either of the bat ANP32 proteins.

**Figure 1.**
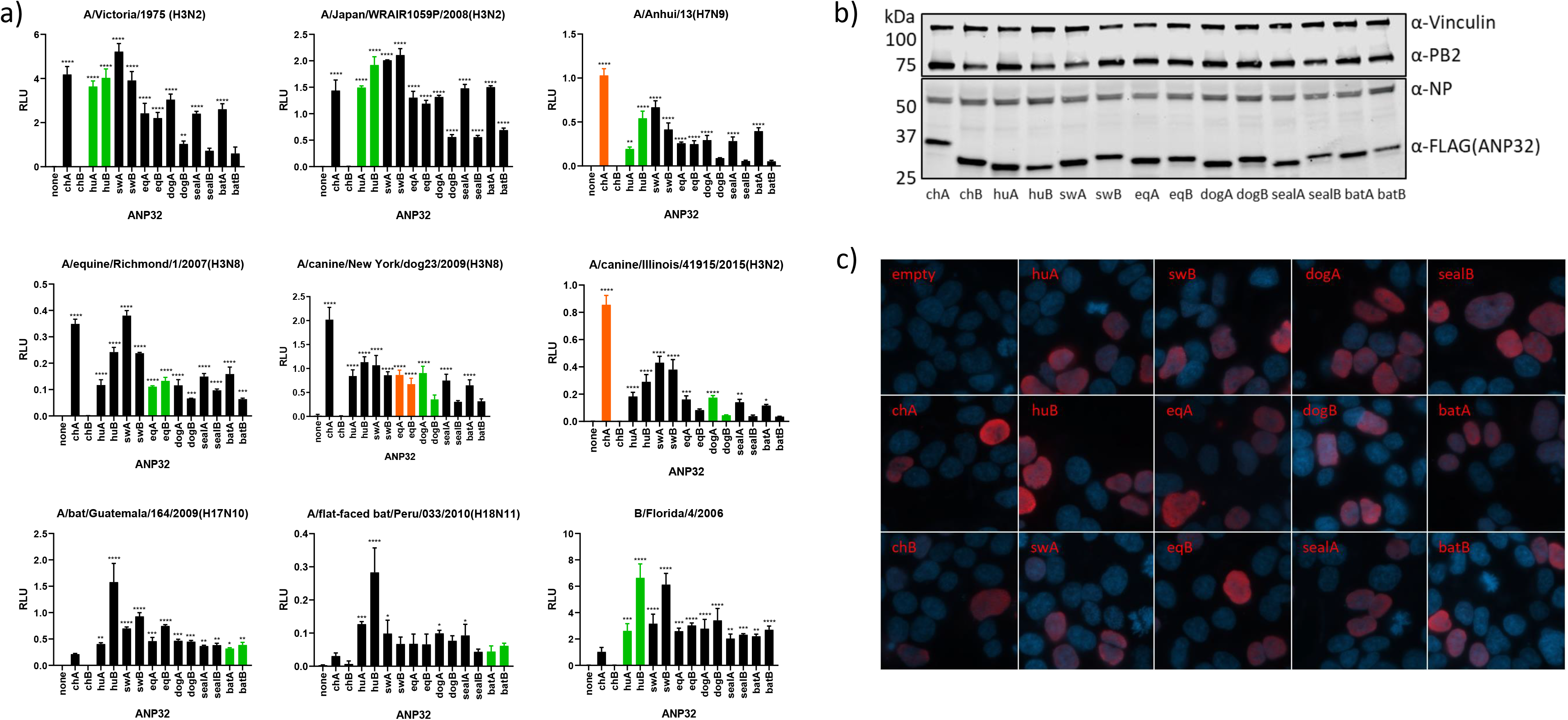
Most mammalian species have two ANP32 proteins capable of supporting influenza polymerase. a) Minigenome assays performed in human eHAP dKO with ANP32 proteins from different avian or mammalian species co-transfected. Green bars indicate species the influenza virus polymerase was isolated from, orange bars indicate recent species the virus has jumped from. b) Western blot assay showing protein expression levels of FLAG-tagged ANP32 proteins during a minigenome assay. c) Immunofluorescence images showing nuclear localisation of all FLAG-tagged ANP32 proteins tested. Abbreviations: ch – chicken, hu – human, sw – swine, eq – equine. Statistical significance was determined by one-way ANOVA with multiple comparisons against empty vector. *, 0.05 ≥ P > 0.01; **, 0.01 ≥ P > 0.001; ***, 0.001 ≥ P > 0.0001; ****, P ≤ 0.0001.

We next tested the ANP32 preference of a human 2009 (swine-origin) pH1N1 and two polymerases from swine influenza isolates. Interestingly, these polymerases were robustly supported by chicken and swine ANP32A, but not other mammalian ANP32 proteins, with the Eurasian avian-like polymerase from A/swine/England/453/2006 (H1N1; sw/453) showing the clearest effect (Fig. 2a). We went on to test a panel of avian-origin viral polymerases with no known mammalian polymerase adaptations, including A/duck/Bavaria/77(H1N1), thought to be an avian precursor of the Eurasian avian-like swine lineage (Fig. 2b). For all the avian origin viral polymerases the stringent preference for avian ANP32A to support polymerase activity was evident (co-expression of chicken ANP32A led to very strong polymerase activity). However, amongst all the mammalian ANP32 proteins tested, only swine ANP32A was able to significantly support avian influenza polymerase activity, though to a lesser degree than chicken ANP32A (Fig. 2b). This unique pro-viral effect of swine ANP32A on swine and avian-origin polymerases was maintained across a wide titration of plasmid doses (Fig. 2c).

**Figure 2.**
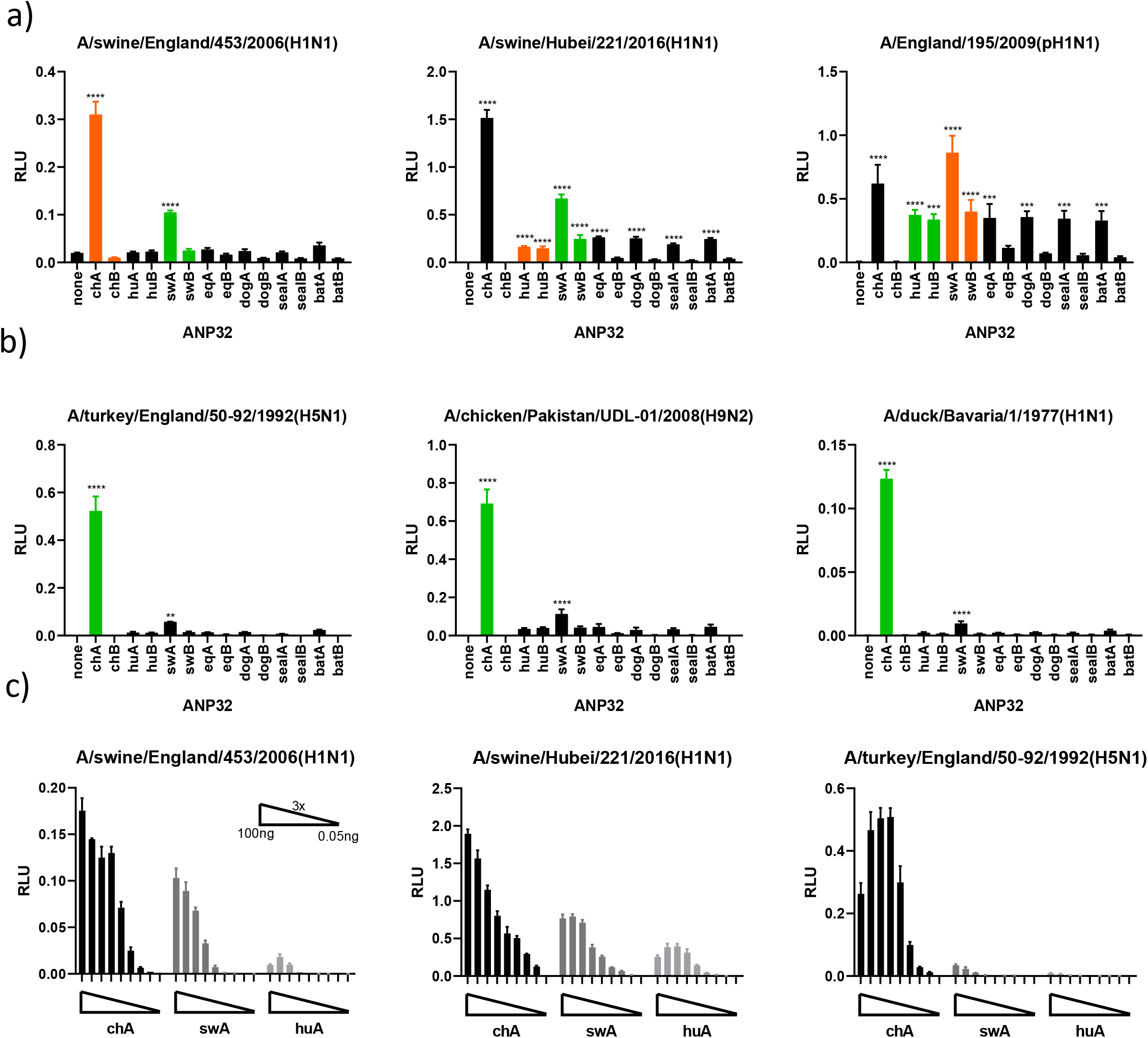
swANP32A can support the activity of minimally mammalian-adapted or completely unadapted polymerases. Minigenome assays of swine (a) and avian (b) polymerases performed in human eHAP dKO cells with ANP32 proteins from different avian or mammalian species co-transfected. Green bars indicate species the influenza virus polymerase was isolated from, orange bars indicate recent species the virus has jumped form. c) ANP32 protein titrations with several different virus polymerases. ANP32 proteins were diluted in a series of 3x dilutions starting with 100ng. Statistical significance was determined by one-way ANOVA with multiple comparisons against empty vector. **, 0.01 ≥ P > 0.001; ***, 0.001 ≥ P > 0.0001; ****, P ≤ 0.0001.

### Swine influenza virus polymerases, adapting to humans, evolve to better use human ANP32 proteins

In 2009 the swine-origin pH1N1 influenza virus adapted from pigs for transmission between humans causing an influenza pandemic (7). The pH1N1 polymerase genes were derived from a triple reassortant constellation in which PB2 and PA originally derived from avian influenza viruses in the mid-1990s (17). From 2009 to 2010 the virus continued to circulate and adapt to humans in the second and third pandemic waves (18). We previously showed that a single substitution in the PA subunit of the polymerase, N321K, contributed to increased polymerase activity of third-wave pH1N1 viruses in human cells (18). We hypothesised that this PA mutation might function by improving support for the emerging virus polymerase by the human ANP32 proteins.

We performed minigenome assays with a first-wave pandemic virus, A/England/195/2009(pH1N1; E195), and a third-wave pandemic virus A/England/687/2010(pH1N1; E687), which differ in PA at position 321. As shown before, PA 321K enhances polymerase activity in general in both virus polymerase backgrounds in human eHAP cells, as well as swine NPTr cells (Fig. 3a). However, the boost is far greater in the human cells (~8-fold) than in the swine cells (~2-fold), implying this mutation may have arisen to overcome the greater restriction seen upon the jump into humans (18).

**Figure 3.**
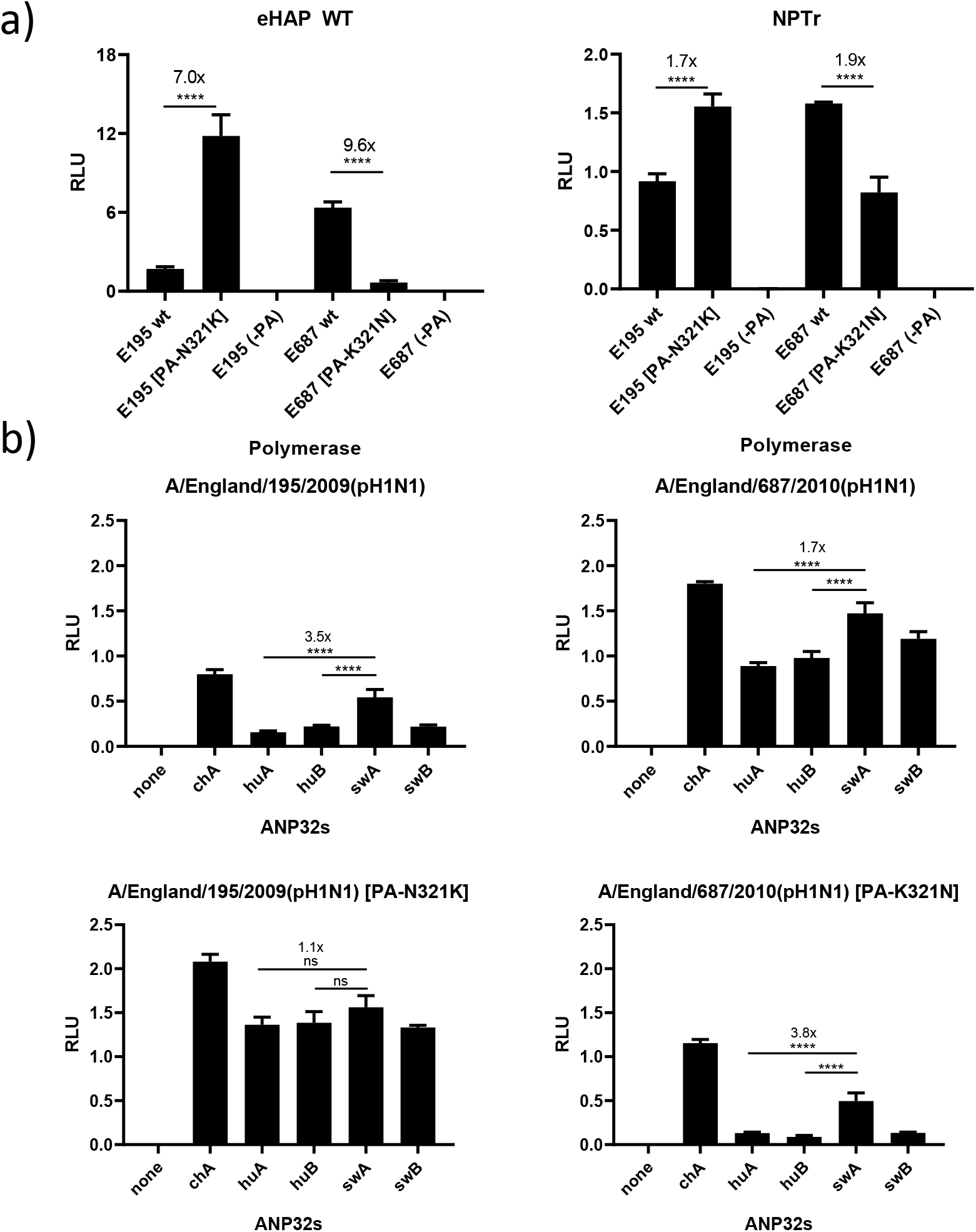
Third-wave pandemic H1N1 viruses adapt to human ANP32 proteins through the PA mutation N321K. a) Minigenome assays of first- and third-wave pH1N1 viruses (E195 and E687, respectively) performed in wild-type human eHAP cells and swine NPTr cells. b) Minigenome assays performed in human eHAP cells with ANP32A and ANP32B knocked out with ANP32 proteins from human or swine co-transfected in. Statistical significance was determined by one-way ANOVA with multiple comparisons. ****, P ≤ 0.0001.

We next tested the ability of human and swine ANP32 proteins to support the different pH1N1 polymerases in eHAP dKO cells. Polymerases containing PA-321N are more robustly enhanced by swine ANP32A (by around 3.5-fold compared to human ANP32A), as is typical of swine-origin polymerases (Fig. 3b). Swine ANP32A, however, gives a much more modest boost to polymerase activity compared to human ANP32A when 321K is present (<2-fold). This suggests the PA N321K adaptation was selected in these viruses to adapt to the more poorly supportive ANP32 proteins present in human cells.

### Differences in swine and human ANP32A pro-viral activity can be mapped to the LRR4 and central region

We set out to identify the molecular basis for the unusually high activity of swine ANP32A in comparison with the other mammalian ANP32 proteins. An alignment of ANP32A primary sequences identified three amino acids outside the LCAR, that differed between swine ANP32A and the other mammalian orthologues. Using reciprocal mutant ANP32A proteins, the identity of amino acid position 156, naturally a serine in swine ANP32A but a proline in most other mammalian ANP32A proteins, was shown to have a major, reciprocal influence on activity (Fig. 4a). The amino acid at position 106 contributed to a lesser degree, with swine-like valine enhancing pro-viral activity over human-like isoleucine. Position 228, localised nearby the C-terminal nuclear localisation signal of ANP32A, had no appreciable impact. In the background of human ANP32A, I106V generally gave between a 1.5- and 6-fold increase in polymerase activity while P156S gave between a 3- and 16-fold boost, depending on the polymerase constellation tested. The combined 106/156 mutant showed an additive effect implying these two residues are, together, responsible for the enhanced pro-viral activity of swine ANP32A (Fig. 4a). None of the mutations affected expression levels (Fig. 4b). Positions 106 and 156 map to the LRR4 and central domains of ANP32 protein, respectively, proximal to the previously characterised LRR5 residues, 129/130, that are responsible for the lack of pro-viral activity of avian ANP32B proteins (Fig. 4c). This reinforces the concept that the LRR4/LRR5/central region of ANP32 proteins is essential to their pro-viral function. Indeed, we could show that introducing the mutation N129I into swine ANP32A abrogated its ability to support influenza polymerase activity (Fig. 4a).

**Figure 4.**
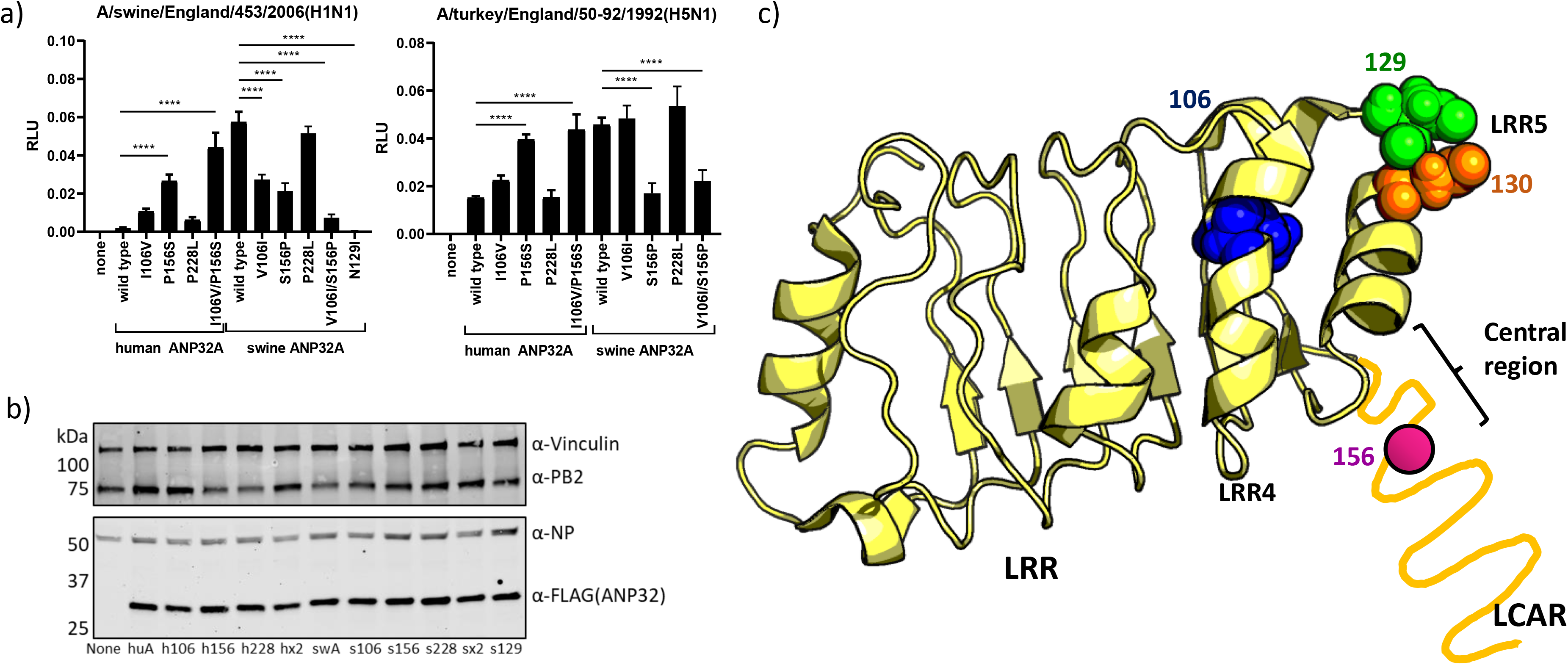
The pro-avian activity of swine ANP32A can be mapped to amino acids in LRR4 and the central domain. a) Minigenome assays performed in human eHAP dKO cells with human/swine ANP32A reciprocal mutants expressed. b) Western blot analysis showing expression levels of human/swine ANP32A from minigenome assays. c) Crystal structure of ANP32 (PDBID: 2JE1) with residues found to affect pro-viral activity mapped (33). The unresolved, unstructured LCAR shown as a yellow line. Schematic made using PyMol (34). Statistical significance was determined by one-way ANOVA with multiple comparisons. *, 0.05 ≥ P > 0.01; ***, 0.001 ≥ P > 0.0001; ****, P ≤ 0.0001.

### An increase in binding to the polymerase accounts for the super pro-viral activity of swine ANP32A

Pro-viral ANP32 proteins from birds and mammals directly bind trimeric polymerase in the cell nucleus (13, 19, 20). Moreover, the inability of avian ANP32B to support influenza polymerase activity correlates with a lack of protein interaction conferred by amino acid differences at residues 129 and 130 (11).

To assess the strength of interaction between swine ANP32A protein and influenza polymerase, we used a split-luciferase assay, where the two halves of *Gaussia* luciferase are fused onto PB1 and ANP32 protein (11, 20). Swine ANP32A interacted strongly with both human-origin - E195 (pH1N1 2009) - and avian-origin - A/turkey/England/50-92/1992(H5N1) - influenza polymerases, although not as strongly as chicken ANP32A (Fig. 5a). Furthermore, the two residues identified as being responsible for strong pro-viral activity of swine ANP32A, at positions 106 and 156, conferred this stronger polymerase binding, implying the mode of action of these mutations is through enhancing ANP32-polymerase interactions (Fig. 5a). It was also shown that N129I, the substitution naturally identified in chicken ANP32B and previously shown to abolish binding and activity in chicken and human ANP32 proteins (11, 15), showed a similar phenotype in swine ANP32A, abolishing detectable binding and activity (Fig. 5a,b). The ablations of the pro-viral activity of swine ANP32A and ANP32B by the substitution N129I was not explained by reductions in expression of these mutant proteins (Fig. 5b,c).

**Figure 5.**
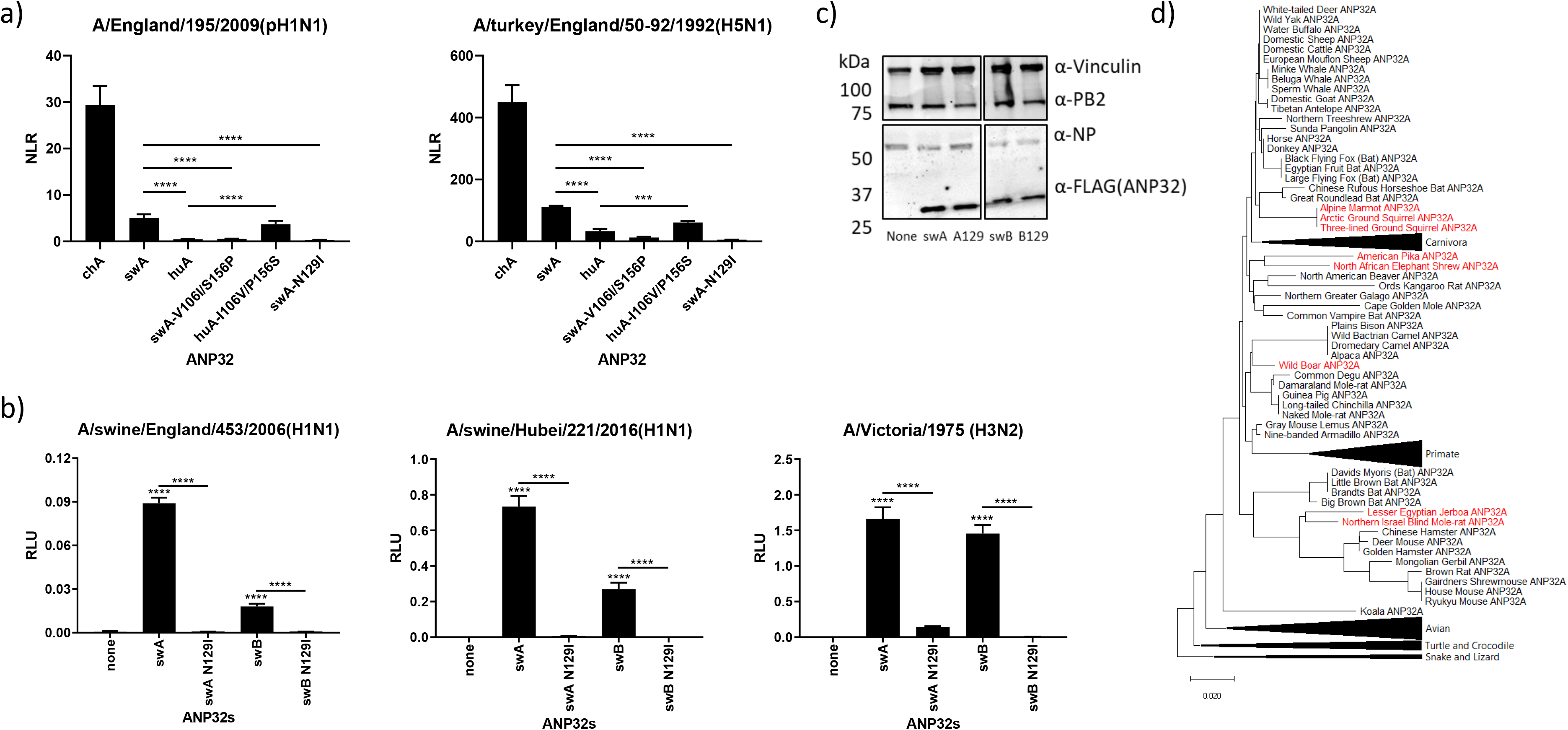
Amino acid residues responsible for the pro-avian polymerase activity of swine ANP32A are also responsible for it binding more strongly to influenza trimeric polymerase. a) Split luciferase assays showing the relative binding of different ANP32 proteins to trimeric polymerase from different influenza virus strains. PB1 was tagged with the N-terminal part of *Gaussia* luciferase while ANP32 proteins were tagged with the C-terminal part. NLR, normalised luminescence ratio, calculated from the ratio between tagged and untagged ANP32/PB1 pairs. Assay performed in 293T cells. Statistical significance was determined by one-way ANOVA with multiple comparisons between the swA and huA wild-types and mutants. ***, 0.001 ≥ P > 0.0001; ****, P ≤ 0.0001. b) Minigenome assays performed in human eHAP cells with ANP32A and ANP32B knocked out with different ANP32 mutants expressed. The N129I naturally occurs in chicken ANP32B. c) Western blot assay showing protein expression levels of FLAG-tagged ANP32 proteins during a minigenome assay. d) phylogenetic tree of mammalian ANP32A proteins, species which contain the highly pro-viral 156S shown in green, species with 156P shown in black. Phylogenetic trees made using the neighbour-joining method based on amino acid sequence. Statistical significance was determined by one-way ANOVA with multiple comparisons against empty vector. ****, P ≤ 0.0001.

### Estimating the pro-viral activity of other mammalian species ANP32A proteins

Based on the molecular markers described in this study it is possible to survey ANP32A proteins from all mammals to predict which other species may have highly influenza polymerase supportive proteins and therefore potential to act as mixing vessels for reassortment between avian and mammalian-adapted influenza viruses.

Very few mammals share the pro-viral marker, 156S, and the few that do mostly constitute species not yet described as hosts for influenza viruses (Fig. 5d). A notable exception is the pika which, in a similar manner to pigs, appears to be highly susceptible to avian influenza viruses with minimal mammalian adaptation (21–23). Pigs are currently the only known mammalian species that contain the secondary, minor pro-viral maker 106V.

## Discussion

In this study we describe the ability of different mammalian ANP32A and ANP32B proteins to support activity of influenza virus polymerases isolated from a variety of hosts. We found that swine ANP32A, uniquely among the ANP32 proteins, supports avian influenza virus polymerase activity. Swine ANP32A does not harbour the avian-specific 33 amino acid duplication that enables the strong interaction and efficient support of polymerase activity of avian-origin viruses by avian ANP32A proteins. Thus, avian influenza viruses are restricted for replication in swine as we have previously shown, and mammalian-adapting mutations enhance their polymerase activity in pig cells (24). Nonetheless, this level of pro-viral activity associated with swine ANP32A, albeit weak compared to avian ANP32As, may contribute to the role of swine as mixing vessels: non-adapted avian influenza viruses that infect pigs could replicate sufficiently to accumulate further mutations that allow for more efficient mammalian adaptation and/or reassortment, enabling virus to either become endemic in swine or to jump into other mammals, including humans.

We map this strongly pro-viral polymerase phenotype to a pair of mutations which allow swine ANP32A to bind more strongly to influenza virus polymerase, potentially explaining the mechanism behind its super pro-viral activity. These residues are only found in a handful of other mammals including wild pigs and pika. It is conceivable these residues are located at a binding interface between polymerase and ANP32, but resolution of the structure of the host:virus complex will be required to confirm this hypothesis.

It has long been speculated that swine play a role as ‘mixing vessels’, by acting as host to both human- and avian-origin influenza viruses. This trait may be partially attributed to receptor patterns in swine allowing viruses that bind to both α2,3 linked (i.e. avian-like viruses) and α2,6 linked sialic acid (i.e. human-like) to replicate alongside each other (8, 9). However, replication of the avian-origin influenza virus genomes inside infected cells is also required to enhance the opportunity for further adaptation or reassortment. We previously developed a minigenome assay for assessing polymerase activity in swine cells and showed that avian virus polymerases were restricted and that restriction could be overcome by typical mutations known to adapt polymerase to human cells (24). Taken together the ability to enter swine cells without receptor switching changes in the haemagglutinin gene, along with a greater mutation landscape afforded in swine cells by the partially supportive pro-viral function of swine ANP32A may have an additive effect to allow swine to act an intermediate host for influenza viruses to adapt to mammals. Furthermore, our work implies other mammals, such as the pika, could play a similar role which is of particular interest due to the pika’s natural habitat often overlapping with that of wild birds and its (somewhat swine-like) distribution of both α2,3 and α2,6-linked sialic acid receptors (25).

Upon crossing into humans from swine, it is likely that viruses would be under selective pressure to adapt to human pro-viral factors, such as the ANP32 proteins. We use the example of a pair of first- and third-wave pandemic H1N1 influenza viruses isolated from clinical cases in 2009 and 2010 (18). The polymerase constellation of the 2009 pH1N1 virus contains PB2 and PA gene segments donated from avian sources to a swine virus in a triple reassortant constellation in the mid-1990s, then passed onto humans in 2009 (17). Although the first-wave viruses, derived directly from swine, can clearly replicate and transmit between humans, over time the PA substitution, N321K, was selected because it enabled more efficient support of the viral polymerase by human ANP32 proteins – our data suggests this is a direct adaptation to the less supportive human ANP32 proteins. This again illustrates how swine have acted as a ‘halfway house’ for the step-wise adaptation of genes originating in avian influenza viruses that have eventually become humanised.

Also of note, we show here that as for the human orthologues, the ANP32A and B proteins of swine (as well as all other mammals tested here) are fairly redundant in their pro influenza activity to support the viral polymerase. We further show that the substitution N129I is able to partially or fully ablate the pro-viral activity of both swine ANP32A and ANP32B. We suggest that the introduction of this substitution in both swine ANP32A and ANP32B by genome editing would be a feasible basis generating influenza resistant, or resilient, pigs, in a similar manner to that demonstrated for porcine respiratory and reproductive syndrome virus resistant pigs, and proposed for influenza resistant, or resilient, chickens (11, 26).

To conclude, we hypothesise that the superior pro-viral function of swine ANP32A for supporting influenza replication may play a role in both the ability of swine to host avian influenza viruses, but also upon the potential evolutionary ecology of swine influenza viruses. This, in turn, may influence the ability of different swine influenza viruses to act as zoonotic hosts or as potential pandemic viruses.

## Materials and methods

### Cells

Human engineered-Haploid cells (eHAP; Horizon Discovery) and eHAP cells with ANP32A and ANP32B knocked out (dKO) by CRISPR-Cas9, as originally described in (14), were maintained in Iscove’s Modified Dulbecco’s Medium (IMDM; ThermoFisher) supplemented with 10% fetal bovine serum (FBS; Biosera), 1% non-essential amino acids (NEAA; Gibco) and 1% Penicillin-streptomycin (pen-strep; invitrogen). Human embryonic kidney (293Ts, ATCC) and Newborn Pig Trachea cells (NPTr; ATCC) were maintained in Dulbecco’s Modified Eagle Medium (DMEM) supplemented with 10% FBS, 1% NEAA and 1% pen-strep. All cells were maintained at 37°C, 5% CO_2_.

### ANP32 plasmids constructs

Animal ANP32 constructs were codon optimised and synthesised by GeneArt (ThermoFisher). Sequences used were pig (*Sus scrofa*) ANP32B (XP_020922136.1), Horse (*Equus caballus*) ANP32A (XP_001495860.2) and ANP32B (XP_023485491.1), Dog (*Canis lupus familiaris*) ANP32A (NP_001003013.2), Dingo (*Canis lupus dingo*) ANP32B (XP_025328134.1), Monk Seal (*Neomonachus schauinslandi*) ANP32A (XP_021549451.1) and ANP32B (XP_021546921.1), and Common Vampire Bat (*Desmodus rotundus*) ANP32A (XP_024423449.1) and ANP32B (XP_024415874.1). All isoforms were chosen due to their equivalence to the known functional human isoforms. Species of origin were chosen due to being flu hosts or the most-commonly related species to flu hosts (in the case of Monk Seal which are closely related to Harbour Seal whereas common vampire bats belong to the same family as little yellow-shouldered and flat-faced bats). Dingo ANP32B was substituted for dog ANP32B as the equivalent isoform used for all other ANP32Bs is unannotated in the dog genome due to a gap in the scaffold. All ANP32 expression constructs included a C-terminal GSG-linker followed by a FLAG tag and a pair of stop codons. Overlap extension PCR was used to introduce mutations into the ANP32 constructs which were then subcloned back into pCAGGS and confirmed by sanger sequencing.

### Viral minigenome plasmid constructs

Viruses and virus minigenome full strain names used through this study were A/Victoria/1975(H3N2; Victoria), A/England/195/2009(pH1N1; E195), A/England/687/2010(pH1N1; E687), A/Japan/WRAIR1059P/2008(H3N2; Japan), B/Florida/4/2006 (B/Florida), A/Anhui/2013(H7N9; Anhui), A/duck/Bavaria/1/1977(H1N1, Bavaria), A/turkey/England/50-92/1992(H5N1; 50-92), A/chicken/Pakistan/UDL-01/2008(H9N2; UDL1/08), A/canine/New York/dog23/2009(H3N8; CIV H3N8), A/canine/Illinois/41915/2015(H3N2; CIV H3N2), A/equine/Richmond/1/2007(H3N8; Richmond), A/swine/England/453/2006(EAH1N1; sw/453), A/swine/Hubei/221/2016(H1N1; Hubei), A/little yellow-shouldered bat/Guatemala/164/2009(H17N10; H17) and A/flat-faced bat/Peru/033/2010(H18N11; H18). Viral minigenome expression plasmids (for PB2, PB1, PA and NP) for H3N2 Victoria, H5N1 50-92, H1N1 E195, H1N1 E687 IBV Florida/06, H9N2 UDL1/08 and H1N1 Bavaria have been previously described (10, 18, 24, 27). Viral minigenome plasmids for H1N1 swine/453, H3N2 Japan, H3N2 CIV, H3N8 CIV, Hubei and Richmond were subcloned from reverse genetics plasmids or cDNA into pCAGGS expression vectors using virus segment specific primers.

pCAGGs minigenome reporters for H17N10 and H18N11 bat influenza viruses were a kind gift from Professor Martin Schwemmle, Universitätsklinikum Freiburg (28). pCAGGs minigenome reporters for H7N9 were a kind gift from Professor Munir Iqbal, The Pirbright Institute, UK. Reverse genetics plasmids for H3N8, Richmond were a kind gift from Adam Rash of the Animal Health Trust, Newmarket, UK. Reverse genetics plasmids for H3N2 CIV and H3N8 CIV were a kind gift from Dr. Colin Parrish of the Baker Institute for Animal Health, Cornell University (29, 30). Viral RNA from sw/453 was kindly provided by Dr. Sharon Brookes, Animal Plant and Health Agency, Weybridge, UK.

### Minigenome assay

eHAP dKO cells were transfected in 24 well plates using lipofectamine^®^ 3000 (thermo fisher) with a mixture of plasmids; 100ng of pCAGGs ANP32/empty vector, 40ng of pCAGGs PB2, 40ng of pCAGGs PB1, 20ng of pCAGGs PA, 80ng of pCAGGs NP, 40ng of pCAGGs *Renilla* luciferase, 40ng of polI vRNA-Firefly luciferase. Transfections in wild-type eHap cells were performed similarly but without ANP32. Transfections in NPTr cells were carried out in 12 well plates using the same ratios above. 24 hours post-transfection cells were lysed with passive lysis buffer (Promega) and luciferase bio-luminescent signals were read on a FLUOstar Omega plate reader (BMG Labtech) using the Dual-Luciferase^®^ Reporter Assay System (Promega). Firefly signal was divided by *Renilla* signal to give relative luminescence units (RLU).

### Split Luciferase Assay

Split luciferase assays were undertaken in 293Ts seeded in 24 well plates. 30ng each of PB2, PA, and PB1, with the N-terminus of *Gaussia* Luciferase (Gluc1) tagged to its C-terminus after a GGSGG linker, were co-transfected using lipofectamine 3000 along with ANP32A, tagged with the C-terminus of *Gaussia* Luciferase (Gluc2) on its C-terminus (after a GGSGG linker). 24 hours later cells were lysed in 100μl of *Renilla* lysis buffer (Promega) and *Gaussia* activity was measured using a *Renilla* luciferase kit (Promega) on a FLUOstar Omega plate reader (BMG Labtech). Normalised luminescence ratios (NLR) were calculated by dividing the values of the tagged PB1 and ANP32 wells by the sum of the control wells which contained 1) untagged PB1 and free Gluc1 and 2) untagged ANP32A and free Gluc2 as described elsewhere (11, 31).

### Western Blotting

To confirm equivalent protein expressing during mini-genome assays transfected cells were lysed in RIPA buffer (150mM NaCl, 1% NP-40, 0.5% Sodium deoxycholate, 0.1% SDS, 50mM TRIS, pH 7.4) supplemented with an EDTA-free protease inhibitor cocktail tablet (Roche).

Membranes were probed with mouse α-FLAG (F1804, Sigma), rabbit α-Vinculin (AB129002, Abcam), rabbit α-PB2 (GTX125926, GeneTex) and mouse α-NP ([C43] ab128193, Abcam). The following near infra-red (NIR) fluorescent secondary antibodies were used: IRDye^®^ 680RD Goat Anti-Rabbit (IgG) secondary antibody (Ab216777, Abcam) and IRDye^®^ 800CW Goat Anti-Mouse (IgG) secondary antibody (Ab216772, Abcam). Western Blots were visualised using an Odyssey Imaging System (LI-COR Biosciences).

### Immunofluorescence

For investigating localisation of exogenously expressed ANP32 proteins, eHAP ANP32 dKO cells were cultured on 8 well chambered cover slips (Ibidi) and transfected with 125 ng of the indicated FLAG-tagged ANP32 protein. Cells were fixed in PBS, 4% paraformaldehyde 24 hours post transfection, then permeabilised in PBS, 0.2% Triton X-100. Cells were blocked in PBS, 2% bovine serum albumin and 0.1% tween. FLAG-tagged ANP32 proteins were detected using mouse anti-FLAG M2 primary antibody (Sigma), followed by goat anti-mouse Alexa Fluor 568 (Invitrogen). Nuclei were counterstained with DAPI. Images were obtained using a Zeiss Cell Observer widefield microscope with ZEN Blue software, using a Plan-Apochromat 63x 1.40-numerical aperture oil objective (Zeiss) and processed using FIJI software (32).

## Acknowledgements

The authors would like to thank members of the Barclay lab, as well as Efstathios Giotis of Imperial College for their scientific input and advise for this project.

T.P.P. was supported by BBSRC grant BB/R013071/1; O.C.S. and P.B.L. were supported by Wellcome trust studentships; E.S. was supported by an Imperial College President’s Scholarship; D.H.G. and W.S.B were supported by Wellcome Trust grant 205100; H.Z. was supported by National Natural Science Foundation of China grant, 31761133005; J.S.L. and W.S.B were supported by BBSRC grant BB/K002465/1; W.S.B was supported by BBSRC grant BB/S008292/1.

